# Edge controllability is associated with treatment response to repetitive transcranial magnetic stimulation in depression

**DOI:** 10.64898/2026.07.11.737986

**Authors:** Subham Dey

**Affiliations:** Computational Neurostimulation Research Program; Noninvasive Neuromodulation Unit; Experimental Therapeutics and Pathophysiology Branch; National Institute of Mental Health; Bethesda; MD; USA

**Keywords:** Edge average controllability, Edge modal controllability, Transcranial magnetic stimulation, Diffusion Weighted Imaging

## Abstract

Repetitive transcranial magnetic stimulation (rTMS) is an established treatment for major depressive disorder (MDD), yet variability in treatment response remains a significant challenge. Network control theory provides a framework to quantify how brain networks facilitate state transitions, but prior work has focused primarily on node level metrics. Here, I investigate whether edge based controllability of the structural connectome is associated with rTMS outcomes. Twenty five patients with treatment-resistant depression underwent diffusion MRI prior to a five week course of high frequency rTMS targeting the dorsolateral prefrontal cortex. Structural connectomes were constructed using MRtrix3 and the Destrieux atlas, and edge based controllability metrics were computed at baseline. Controllability of specific middle frontal gyrus centered edges showed significant associations with changes in HAMD-24 scores, including connections to the superior frontal gyrus, hippocampus, angular gyrus, and orbital gyrus (r = -(0.470–0.597), p < 0.05). These findings suggest that edge based controllability captures circuit level properties relevant to treatment response and may inform personalized neuromodulation strategies.

## I. Introduction

Depression, also known as major depressive disorder, is a mood disorder which affects cognition, behavior, and somatic processes, typically disrupting daily functioning and, in severe instances, causing worthlessness or hopelessness. It is very prevalent, and the World Health Organization (WHO) has estimated that more than 280 million people are affected globally. In the United States alone, nearly 1 in 5 adults experience some form of mental illness every year, with depression being one of the leading causes of disability-adjusted life years [1]. Neuromodulation techniques have shown to be effective therapies for such disorders, supplementing or even in certain cases replacing pharmacologic and psychotherapeutic interventions. Of the noninvasive techniques, transcranial magnetic stimulation (TMS) has been extensively researched and clinically used. The first application of TMS was reported by Barker et al. [2], where a single magnetic pulse applied over the motor cortex induced movement in the contralateral limb. Advances in technology enabling multiple pulses gave rise to repetitive TMS (rTMS) [3], while the neuroimaging revolution allowed more precise targeting, particularly of the left dorsolateral prefrontal cortex (dlPFC). Stimulation frequency was found to be a critical factor: high-frequency rTMS (up to 20 Hz) increases cortical excitability, whereas low-frequency rTMS (around 1 Hz) decreases it [4], [5]. Numerous controlled trials led to FDA clearance of rTMS for depression in 2008, and subsequent advances introduced theta-burst protocols, including accelerated variants, with approvals in 2018 and 2020 [6]–[8].

Network control theory is a field which has its roots in the domains of network systems [9] and control theory [10]. Network control theory hypothesizes the brain as a dynamical system, wherein the concepts of average and modal controllability give us an idea of how easily state transitions happen through the lens of energy landscape [11]. A node is said to have a high value of average controllability if stimulating the node leads to low energy transition whereas a node is said to have a high value of modal controllability if stimulating the node leads to high energy transition. Concepts from network control theory has been previously used as a therapeutic planning mechanism when the treatment mechanism is electroconvulsive therapy (ECT) [12]. ECT is one of the most prominent technique for treating depression. However, most of the works in the literature have considered the controllability metrics from a ‘node’ perspective. To alleviate the limitations related to ‘node’ perspective which includes precision, in this paper the controllability metrics are defined from an ‘edge’ perspective. Most of the work related to ‘edge’ based approaches had focused on functional connectivity rather than on the structural connectivity [13]. In this brief, I shed light on how edge based controllability metrics obtained from the structural connectome can provide an alternative way for associating treatment response for subjects being subjected to rTMS treatment. To my knowledge, this is the first study to investigate edge based controllability of the structural connectome as an associator of rTMS response in depression. The concepts of edge controllability can be extended to morphometric similarity networks as proposed in [14] from the structural connectome concept on which it is proposed in this paper. It can also help us to understand the inherent neurobiological signatures of psychiatric disorders like depression and give us a mechanisitic understanding of how various interventions work to help alleviate the disease symptoms thereby marching towards replicability of intervention paradigms.

The paper is organized in the following subsections, in section II, I explain some of the mathematical preliminaries associated with edge controllability of networked systems. In Section III, some concepts related to the data and preprocessing steps are mentioned. The results of the paper are explained in Section IV whereas the discussions are explained in Section V. The conclusion section of the paper is given in Section VI with the acknowledgement in Section VII.

## II. Mathematical Preliminaries

Taking cue from the work published in [15], here I will state the basics of edge based controllability derived from first principles of network control theory. With inspirations from linear dynamical systems theory on continuous time domain one can represent the evolution of brain states as

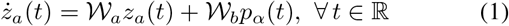

where *W*_*a*_ = [*w*_*i,j*_]_*i,j*=1,2,…, ℕ_ ∈ ℝ^*n*×*n*^ represents the node based adjacency matrix obtained from the structural connectome and N depends on how you parcellate the brain depending on the atlas you choose for parcellation. *z*_*a*_(*t*) ∈ ℝ^*n*×1^ is used to represent the state whereas *p*_*α*_(*t*) ∈ ℝ^*m*×1^ represents the control input. *W*_*b*_ ∈ ℝ^*n*×*m*^ is used to represent how the control input is applied to the brain. The evolution of brain states in a discrete time setting is given by,

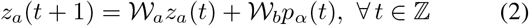

The expression of node average controllability and node modal controllability can be obtained from the expression of controllability gramian in the discrete time setting in which it was postulated in the scientific literature [16].

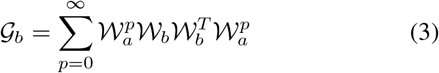

Now in order to have the expression of the average controllability I use the following expression trace(*G*_*b*_) since the expression trace 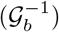 may result in computational ill-posedness (wherein trace of a matrix is the sum of elements on its diagonal) [17].

I construct a matrix *W*_*eg*_ = [*o*_*ij*_] which represents normalized eigenvectors of matrix *W*_*a*_ from the discrete time equation given in Eq. (2) . *o*_*ij*_ reflects the controllability of the mode *λ*_*j*_(*W*_*a*_) provided the control node is the node i. One can express the scaled measure of controllability of all modes {*λ*_*i*_} _*i*=1,2,3,…., ℕ_ from the control node *i* through the aid of Popov Belevitch Hautus (PBH) test as [18]

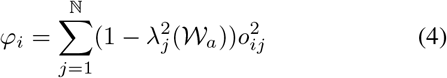

The expression of incidence matrix *C*_*a*_ can be obtained through the aid of the following equation

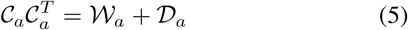

where *D*_*a*_ = diag(*f*_*ii*_)_*i*=1,2,…,N_ and *f*_*ii*_ = ∑_*j*_ (*w*_*ij*_). Now the edge adjacency matrix can be written as

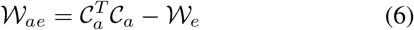

where *W*_*e*_ represents a diagonal matrix with its elements representing the weight of each edge. The discrete time state space representation of the neural dynamics in the edge based formulation is given by

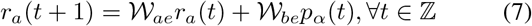

where *W*_*ae*_ ∈ ℝ ^(0.5∗*n*∗(*n*−1))×(0.5∗*n*∗(*n*−1))^, *r*_*a*_(*t*) ∈ ℝ ^(0.5∗*n*∗(*n*−1))^ represents the edge state variable, *p*_*α*_ ∈ ℝ^*r*×1^ represents the control input and finally *W*_*be*_ ∈ ℝ ^(0.5∗*n*∗*n*−1)×*r*^ represents how control input is applied to the edge graph. Edge average controllability can be obtained with the help of a gramian matrix through the aid of control matrix _*be*_. The expression of the corresponding gramian matrix is given by

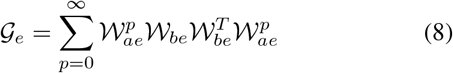

The expression of edge average controllability is given by trace(*G*_*e*_) (wherein trace of a matrix is the sum of elements on its diagonal). Similarly, the expression of edge modal controllability is given by

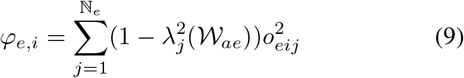

*o*_*eij*_ represents the normalized eigenvectors of the edge adjacency matrix. The variable ℕ_*e*_ reflects the dimensions of edge adjacency matrix under consideration.

## III. Data and preprocessing

The Institutional Review Board (IRB) of Weill Cornell Medical College approved this study, and all participants provided written informed consent prior to enrollment. Twenty five currently treatment-resistant depressed patients (age 21-68) received daily 10-Hz rTMS over dlPFC for five days per week for five weeks. Treatment response was assessed using the 24-item Hamilton Rating Scale for Depression (HAMD), administered at baseline and following completion of the treatment course. Single shell diffusion tensor images at 3T were acquired within 7 days prior to and within 3 days after the rTMS treatment course. A high-resolution T1-weighted anatomical volume was also acquired for each subject to support tissue segmentation and registration.

Diffusion weighted imaging data were processed using MR-trix3 [19] in combination with FSL [20] and FreeSurfer tools [21]. Preprocessing began with denoising via Marchenko-Pastur principal component analysis (`dwidenoise`), followed by removal of Gibbs ringing artifacts. Correction for eddy current distortions, subject motion, and susceptibility induced geometric distortion was performed using FSL’s `eddy`, and bias field correction was applied using the ANTs based N4 algorithm implemented in MRtrix3. A brain mask was generated from the preprocessed diffusion data and refined using morphological operations to exclude non-brain tissue. Fiber orientation distributions (FODs) were estimated using single-shell constrained spherical deconvolution (CSD) with the Tournier algorithm for white matter response function estimation, using a maximum spherical harmonic order, followed by multi-tissue intensity normalization to correct for global intensity differences across subjects.

Structural T1 weighted images were processed using FreeSurfer’s standard `recon-all` pipeline to obtain cortical surface reconstructions and anatomical segmentations, including parcellation according to the Destrieux atlas. Diffusion images were registered to anatomical space using linear registration (FLIRT [22], 6 degrees of freedom), and the resulting transformation was applied to align tissue segmentations and parcellations to diffusion space; registration quality was visually inspected for each subject.

Whole-brain probabilistic tractography was performed with anatomically constrained tractography (ACT), using the segmented tissue boundaries to restrict streamline propagation to biologically plausible pathways. Streamlines were seeded from the gray matter–white matter interface using dynamic seeding, with a total of 10 million streamlines generated per subject, with a predetermined maximum curvature angle and a step size. To improve biological accuracy and correct for known biases in streamline density, streamline weights were adjusted using the SIFT2 algorithm [23]. Structural connectomes were constructed using the Destrieux atlas (164 regions) [24], with edges defined as the SIFT2 weighted sum of streamlines connecting each pair of regions. Connectomes were symmetrized, diagonal elements were excluded, and edge weights were normalized by inverse node volume to account for variability in regional size across subjects. The edge connectome will be of dimension 13366 × 13366 using the formulae 0.5\**n\**(*n* − 1) where n is the dimension of structural connectome.

## IV. Results

Edge average controllability and edge modal controllability values were calculated from the edge based structural connectome obtained at the baseline. Restricting the analysis to edges which are consistently present across all subjects as well as those in which one node of the edge is either left or right middle frontal gyrus parcellation of the Destrieux atlas I found that a subset of edges showed significant associations with changes in HAMD scores (*p <* 0.05). The list of edges are given in Table 1. The plots related to association between changes in HAMD with respect to edge average controllability are given in Figs. 1 and 2 respectively. The plots also contains people those who responded to treatment and those who are non-responders colored in different colors (non-responder blue and responder orange) where responders refer to people whose HAMD scores decreases greater than 50 % with respect to baseline values after rTMS treatment. Whereas nonresponders are those whose HAMD scores didnot decrease greater than 50 % with respect to baseline values after rTMS treatment. The plots related to edge modal controllability and changes in HAMD scores are not shown as the relationship between average and modal controllability follow an inverse law thereby satisfying the energy landscape hypothesis which is the backbone of the controllability analysis.

**TABLE I.**

| Edge | Pearson correlation (r) | p-value (uncorrected) |
| --- | --- | --- |
| Left middle frontal gyrus–right superior frontal gyrus | -0.597 | 0.00163 |
| Left middle frontal gyrus–left hippocampus | -0.547 | 0.0046 |
| Right middle frontal gyrus–right angular gyrus | -0.525 | 0.0071 |
| Right middle frontal gyrus–right orbital gyrus | -0.470 | 0.01764 |

**Fig. 1.**
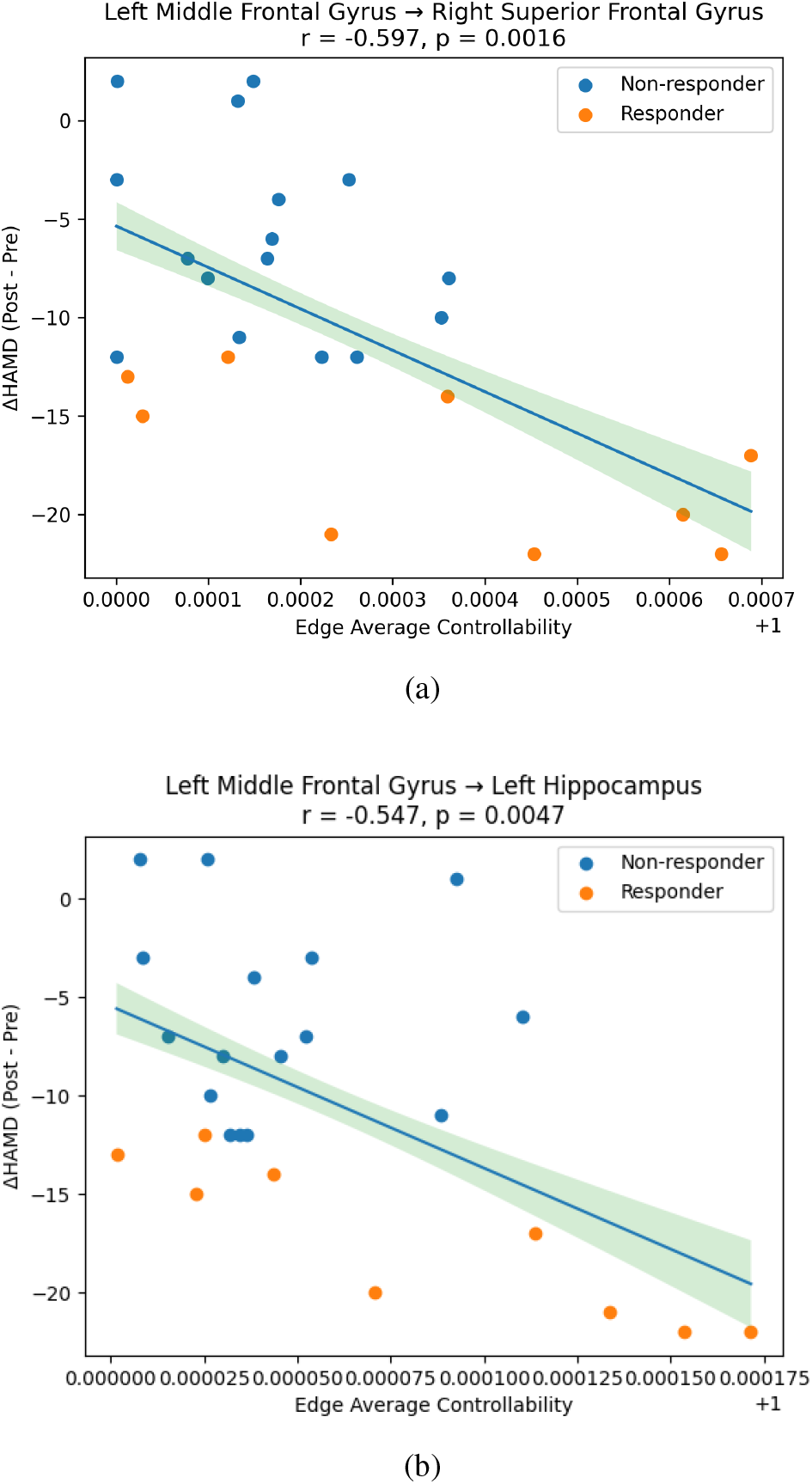
Edge average controllability and relationship between changes in HAMD scores after treatment through rTMS treatment protocol (a) Left middle frontal gyrus–right superior frontal gyrus, (b) Left middle frontal gyrus–left hippocampus.

**Fig. 2.**
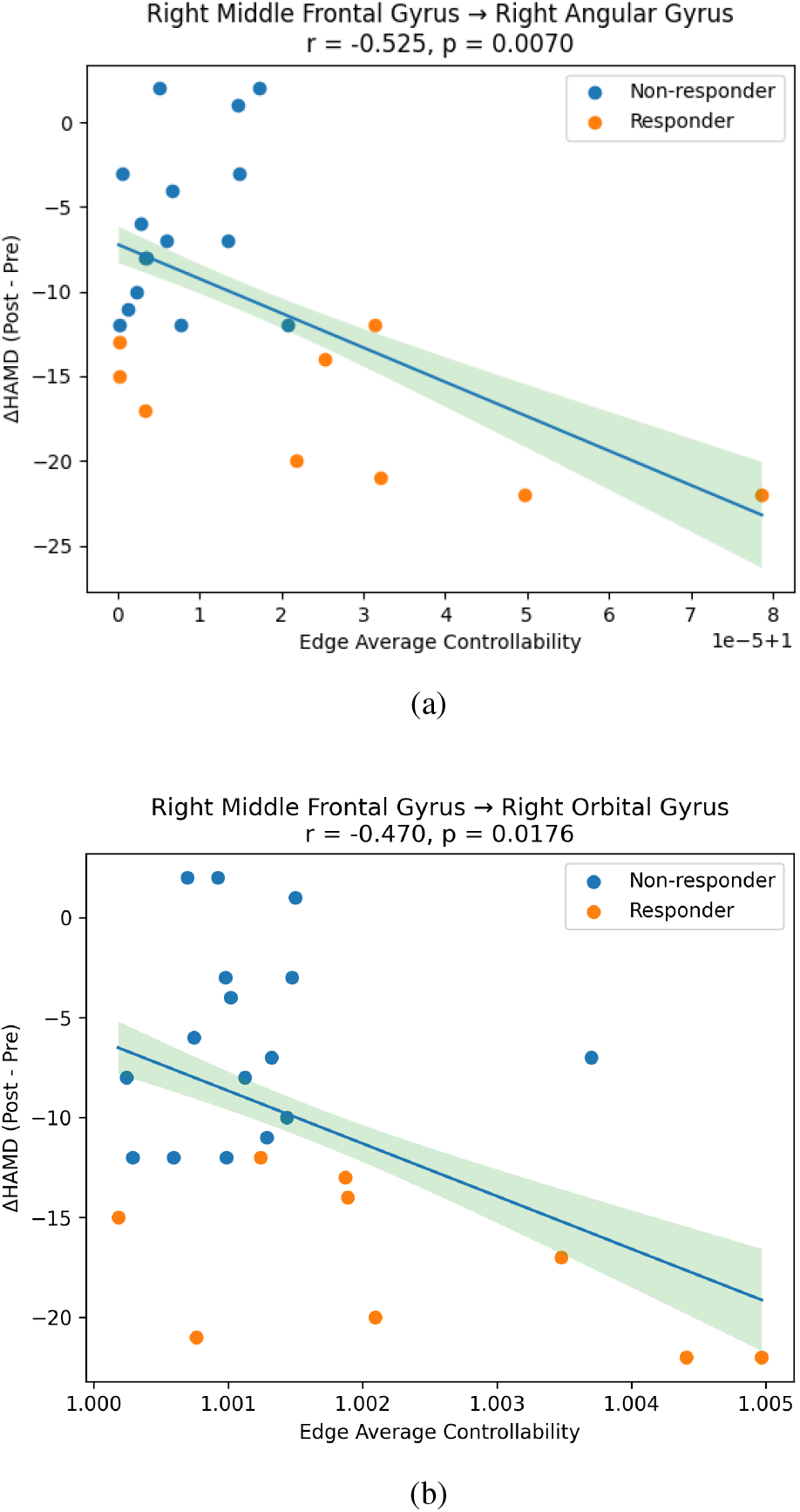
Edge average controllability continued: (a) Right middle frontal gyrus–right angular gyrus, (b) Right middle frontal gyrus–right orbital gyrus.

## V. Discussion

In this work, I elucidate that edge controllability metrics derived from structural connectome at the baseline are related with antidepressant response to repetitive transcranial magnetic stimulation (rTMS) in patients with treatment resistant depression. Significantly, controllability of edges linking middle frontal gyrus to superior frontal gyrus, hippocampus, angular gyrus, and orbital gyrus showed robust correlations with changes in HAMD scores. These findings provide support for the hypothesis that the efficacy of rTMS is not entirely determined by local stimulation targets, but rather by the ability of distributed network connections to facilitate global brain state transitions.

A key finding of this study is the identification of frontolimbic and fronto-associative circuits as predictors of treatment response. In particular, the relationship between middle frontal gyrus and hippocampus connectivity and symptom improvement highlights the role of prefrontal modulation of limbic structures involved in memory and affective processing. Similarly, connections between the middle frontal gyrus and orbitofrontal cortex were associative of treatment outcome, implicating reward and valuation circuits that are frequently disrupted in depression. The involvement of fronto-parietal connections, such as those linking the middle frontal gyrus to the angular gyrus, further supports the importance of large scale cognitive control and default mode networks in mediating therapeutic response. A major contribution of this work is the shift from node based to edge based controllability analysis. While prior studies have primarily focused on node level metrics, such approaches may lack the spatial precision required to capture specific pathways mediating treatment effects. This edge centric perspective offers a more refined understanding of brain network control and may be particularly relevant for guiding targeted neuromodulation strategies.

Several limitations should be considered. First, the sample size is modest (n = 25), which may limit generalizability and statistical power. Second, the analysis is based on structural connectivity derived from diffusion MRI, which is subject to limitations in tractography accuracy and may not fully capture functional or synaptic interactions. Third, the study relies on correlational analysis and does not establish causality between controllability metrics and treatment response.

## VI. Conclusion

This study demonstrates that baseline edge based controllability of the structural connectome is associated with clinical response to rTMS in treatment resistant depression. By identifying specific middle frontal gyrus specific connections whose controllability relates to symptom improvement, our findings provide a network level framework for understanding variability in treatment outcomes.

These results highlight the potential of edge centric metrics as objective markers to inform patient selection and guide neuromodulation strategies. Future work should focus on validating these findings in larger cohorts and evaluating their associative utility in prospective clinical settings.

## VII. Acknowledgment

The author is supported by the National Institute of Mental Health (NIMH) Intramural Research Program (ZIAMH002955). AI tools were used in preparation of the manuscript but the author takes full responsibility for the statements made herein.

## References

[1] P. E. Holtzheimer and H. S. Mayberg, “Stuck in a rut: rethinking depression and its treatment,” Trends in neurosciences, vol. 34, no. 1, pp. 1–9, 2011.

[2] A. T. Barker, R. Jalinous, and I. L. Freeston, “Non-invasive magnetic stimulation of human motor cortex,” The Lancet, vol. 325, no. 8437, pp. 1106–1107, 1985.

[3] G. Höflich, S. Kasper, A. Hufnagel, S. Ruhrmann, and H.-J. Möller, “Application of transcranial magnetic stimulation in treatment of drug-resistant major depression—a report of two cases,” Human Psychopharmacology: Clinical and Experimental, vol. 8, no. 5, pp. 361–365, 1993.

[4] A. Pascual-Leone, J. Valls-Solé, E. M. Wassermann, and M. Hallett, “Responses to rapid-rate transcranial magnetic stimulation of the human motor cortex,” Brain, vol. 117, no. 4, pp. 847–858, 1994.

[5] R. Chen, J. Classen, C. Gerloff, P. Celnik, E. Wassermann, M. Hallett, and L. G. Cohen, “Depression of motor cortex excitability by low-frequency transcranial magnetic stimulation,” Neurology, vol. 48, no. 5, pp. 1398–1403, 1997.

[6] P. E. Holtzheimer and W. McDonald, A clinical guide to transcranial magnetic stimulation. Oxford University Press, 2014.

[7] D. M. Blumberger, F. Vila-Rodriguez, K. E. Thorpe, K. Feffer, Y. Noda, P. Giacobbe, Y. Knyahnytska, S. H. Kennedy, R. W. Lam, Z. J. Daskalakis et al., “Effectiveness of theta burst versus high-frequency repetitive transcranial magnetic stimulation in patients with depression (three-d): a randomised non-inferiority trial,” The Lancet, vol. 391, no. 10131, pp. 1683–1692, 2018.

[8] E. J. Cole, K. H. Stimpson, B. S. Bentzley, M. Gulser, K. Cherian, C. Tischler, R. Nejad, H. Pankow, E. Choi, H. Aaron et al., “Stanford accelerated intelligent neuromodulation therapy for treatment-resistant depression,” American Journal of Psychiatry, vol. 177, no. 8, pp. 716–726, 2020.

[9] Y.-Y. Liu, J.-J. Slotine, and A.-L. Barabási, “Controllability of complex networks,” nature, vol. 473, no. 7346, pp. 167–173, 2011.

[10] F. Pasqualetti, S. Zampieri, and F. Bullo, “Controllability metrics, limitations and algorithms for complex networks,” IEEE Transactions on Control of Network Systems, vol. 1, no. 1, pp. 40–52, 2014.

[11] L. Parkes, J. Z. Kim, J. Stiso, J. K. Brynildsen, M. Cieslak, S. Covitz, R. E. Gur, R. C. Gur, F. Pasqualetti, R. T. Shinohara et al., “A network control theory pipeline for studying the dynamics of the structural connectome,” Nature Protocols, vol. 19, no. 12, pp. 3721–3749, 2024.

[12] T. Hahn, H. Jamalabadi, E. Nozari, N. R. Winter, J. Ernsting, M. Gruber, M. J. Mauritz, P. Grumbach, L. Fisch, R. Leenings et al., “Towards a network control theory of electroconvulsive therapy response,” PNAS nexus, vol. 2, no. 2, p. pgad032, 2023.

[13] M. A. de Reus, V. M. Saenger, R. S. Kahn, and M. P. van den Heuvel, “An edge-centric perspective on the human connectome: link communities in the brain,” Philosophical Transactions of the Royal Society B: Biological Sciences, vol. 369, no. 1653, 2014.

[14] I. Sebenius, J. Seidlitz, V. Warrier, R. A. Bethlehem, A. Alexander-Bloch, T. T. Mallard, R. R. Garcia, E. T. Bullmore, and S. E. Morgan, “Robust estimation of cortical similarity networks from brain mri,” Nature neuroscience, vol. 26, no. 8, pp. 1461–1471, 2023.

[15] S. Dey, E. Bharti, and Z.-D. Deng, “Controllability analysis of macaque structural connectome from an edge centric perspective,” in 2025 47th Annual International Conference of the IEEE Engineering in Medicine and Biology Society (EMBC). IEEE, 2025, pp. 1–6.

[16] S. Gu, F. Pasqualetti, M. Cieslak, Q. K. Telesford, A. B. Yu, A. E. Kahn, J. D. Medaglia, J. M. Vettel, M. B. Miller, S. T. Grafton et al., “Controllability of structural brain networks,” Nature communications, vol. 6, no. 1, p. 8414, 2015.

[17] E. Tang, C. Giusti, G. L. Baum, S. Gu, E. Pollock, A. E. Kahn, D. R. Roalf, T. M. Moore, K. Ruparel, R. C. Gur et al., “Developmental increases in white matter network controllability support a growing diversity of brain dynamics,” Nature communications, vol. 8, no. 1, p. 1252, 2017.

[18] T. Kailath, Linear systems. Prentice-hall Englewood Cliffs, NJ, 1980, vol. 156.

[19] J.-D. Tournier, R. Smith, D. Raffelt, R. Tabbara, T. Dhollander, M. Pietsch, D. Christiaens, B. Jeurissen, C.-H. Yeh, and A. Connelly, “Mrtrix3: A fast, flexible and open software framework for medical image processing and visualisation,” Neuroimage, vol. 202, p. 116137, 2019.

[20] M. Jenkinson, C. F. Beckmann, T. E. Behrens, M. W. Woolrich, and S. M. Smith, “Fsl,” Neuroimage, vol. 62, no. 2, pp. 782–790, 2012.

[21] B. Fischl, “Freesurfer,” Neuroimage, vol. 62, no. 2, pp. 774–781, 2012.

[22] S. M. Smith, M. Jenkinson, M. W. Woolrich, C. F. Beckmann, T. E. Behrens, H. Johansen-Berg, P. R. Bannister, M. De Luca, I. Drobnjak, D. E. Flitney et al., “Advances in functional and structural mr image analysis and implementation as fsl,” Neuroimage, vol. 23, pp. S208–S219, 2004.

[23] R. E. Smith, J.-D. Tournier, F. Calamante, and A. Connelly, “Sift2: Enabling dense quantitative assessment of brain white matter connectivity using streamlines tractography,” Neuroimage, vol. 119, pp. 338–351, 2015.

[24] C. Destrieux, B. Fischl, A. Dale, and E. Halgren, “Automatic parcellation of human cortical gyri and sulci using standard anatomical nomenclature,” Neuroimage, vol. 53, no. 1, pp. 1–15, 2010.

